# A longitudinal study of white matter functional network in mild traumatic brain injury

**DOI:** 10.1101/2020.09.25.313338

**Authors:** Xiaoyan Jia, Xuebin Chang, Lijun Bai, Yulin Wang, Debo Dong, Shuoqiu Gan, Shan Wang, Xuan Li, Xuefei Yang, Yinxiang Sun, Tianhui Li, Feng Xiong, Xuan Niu, Hao Yan

**Author notes:** These authors contributed equally to this study (Xiaoyan Jia, Xuebin Chang, Lijun Bai). Correspondence: Hao Yan, Address: Key laboratory for artificial intelligence and cognitive neuroscience of language, Xi’an International Studies University, Xi’ an 710128, China.

## Abstract

The mild traumatic brain injury (mTBI) results in traumatic axonal injury, which damages the long-distance white matter (WM) connections and thus disrupts the functional connectome of large-scale brain networks that support cognitive function. Patterns of WM structural damage following mTBI were well documented using diffusion tensor imaging, however, the functional organization of WM and its association with grey matter functional networks (GM-FNs) and cognitive assessments remains unknown. The present study adopted resting-state functional magnetic resonance imaging to explore WM functional properties in mTBI patients (113 acute patients, 56 chronic patients, 47 healthy controls (HCs)). Eleven large-scale WM functional networks (WM-FNs) were constructed by the k-means clustering algorithm which carried out in voxel-wise WM functional connectivity (FC). Compared to HCs, acute mTBI patients showed enhanced FC between inferior fronto-occipital fasciculus (IFOF) WM-FN and primary sensorimotor WM-FNs, and cortical primary sensorimotor GM-FNs. And FC between IFOF WM-FN and anterior cerebellar GM-FN was positively correlated with information processing speed. Moreover, all of these WM-FNs abnormalities were returned to the normal level at the chronic stage. Our findings suggest the compensatory mechanism of cognitive deficits in the acute stage and its involvement in facilitating recovery from cognitive deficits in the chronic stage. The convergent damage of the IFOF network highlighted its key role in our understanding of the pathophysiology mechanism of mTBI patients and thus might be regarded as a biomarker in the acute stage and a potential indicator of treatment effect.

## Introduction

Traumatic brain injury (TBI) is a public health challenge of vast proportions but insufficiently recognized. More than 50 million people have a TBI worldwide each year, and it is evaluated that about half of the world’s population will have one or more TBIs in their lifetime [1,2].

The mild TBI (mTBI) accounts for 80%~90% of all TBI cases in both civilian and military populations [3]. In general, mTBI results in traumatic axonal injury [4,5]. Previous studies suggested that the traumatic axonal injury would induce cortical network disconnection and disrupt cortical-subcortical pathways in mTBI [6–8], which further impair information transfer across brain networks [9] and produce cognition deficits [10]. Importantly, the long white matter (WM) tracts connections are particularly vulnerable to the mechanical trauma of TBI [11] and thus disrupt the functional communication of large-scale networks that support cognition [12]. For instance, the inferior fronto-occipital fasciculus (IFOF) is a brain region mediates feed-forward propagation of visual input to anterior frontal regions and was found to have direct top-down modulation of early visual processing [13]. IFOF was documented with reduced WM integrity in mTBI patients [14]. Besides, the corpus callosum and internal capsule are regions where projection and contact fibers are highly concentrated. These two regions are responsible for the transmission of important information such as somatosensory and motor signals, and are the most commonly damaged regions in mTBI [15,16]. And previous study evidenced the cognition deficit might due to the damage of WM connecting the widely distributed grey matter (GM) networks [17], suggested the WM play a crucial role in information communication among large-scale brain networks. Although previous studies revealed the alterations in WM tracts in mTBI patients using diffusion tensor imaging (DTI), functional organization of WM and its association with GM functional networks (GM-FNs) across whole brain remains unknown.

Over the past two-decades, resting-state functional magnetic resonance imaging (rs-fMRI) has been a powerful technique for investigating the functional architecture of human GM [18,19]. Recently, increasing studies have evidenced the existence of functional information in WM, which can also be detected by rs-fMRI [20–22]. For instance, Ding et al. uncovered that resting-state BOLD signals in WM reflect neural coding and information processing [20]. Several studies further indicated that the WM would be activated in response to multiple tasks, such as perceptual and motor tasks [23–25]. Besides, the large-scale functional organization of WM has been built [26,27]. Furthermore, recent studies revealed that the functional connectivity (FC) in WM is related to the underlying pathophysiological mechanism of neuropsychiatric disorders, such as Parkinson’s disease [28], schizophrenia [29], and epilepsy [30]. Together, all previous studies have suggested the involvement of functional network of WM in cognitive and disorder progression. Evidently, the WM lesions widely exist in mTBI and the impact of these lesions on cognition is well documented in previous studies. Therefore, the present study constructed the WM-FNs of mTBI patients, and further investigated the longitudinal neuroimaging trajectories of functional properties of WM functional networks (WM-FNs) and its association with GM-FNs in mTBI patients to deepen our understanding of the pathological mechanism of cognitive deficits in mTBI patients.

The present study aims to (i) construct large-scale WM-FNs by k-means clustering analysis on rs-fMRI data of a large cohort of 113 acute mTBI patients (of which 56 patients followed-up at six months to one year, which were considered as chronic stage patients) and 47 healthy controls (HCs). (ii) investigate the FC within the resulting large-scale WM-FNs and its interaction with the known GM-FNs in both acute and chronic mTBI patients. (iii) estimate the spatial interaction between the resulting WM-FNs and structural WM tracts. (iv) measure the correlations between WM functional disturbances and cognitive and clinical assessments of mTBI patients.

## Material and methods

### Participants

One hundred and thirteen consecutive patients suffering from head trauma were recruited from the local emergency department between August 2016 and May 2018 as the initial population. The inclusion and exclusion criteria of mTBI were based on the World Health Organization’s Collaboration Centre for Neurotrauma Task Force [31]. The detailed inclusion and exclusion criteria of mTBI patients were described in the supplementary material. Fifty-six of mTBI patients in the acute stage were followed up for six months to one year, with repeat MRI and cognitive and clinical assessment. Forty-seven age-, gender-, and education-matched HCs with no history of neurologic impairment or psychiatric disorders were included.

MRI scans of patients with mTBI were initially assessed in acute stage (within 7 days post-injury) and followed up at chronic stage (6 months to 1-year post-injury). Cognitive and clinical assessment (see below) was carried out within 48 hours of MRI scans. All HCs participants also underwent an identical MRI scanning and cognitive and clinical assessment.

All participants were right-handed according to the Edinburgh Handedness Inventory [32]. Written informed consent was obtained from each individual before the experimental procedures. The research procedures have been approved by the Local Institutional Review Board (the First Affiliated Hospital of Xi’an Jiaotong University) and conducted in line with the Declaration of Helsinki.

### Image acquisition

A non-contrast CT scan was performed on all consecutive patients following acute head injury with a 64-row CT scanner (GE, Lightspeed VCT). All participants underwent MRI scanning in a 3.0 T MRI scanner (GE 750) with a 32-channel head coil. The detailed scan parameters were described in the supplementary material.

### Cognitive and clinical assessment

Cognitive assessment included: (a) Trail-Making Test Part A (TMT-A) and WAIS-III Digital Symbol Coding score (DSC), which were used to evaluate cognitive information processing speed [33]; (b) WAIS-III Forward Digit Span (DS) and Backward DS, which were used to measure working memory [34]; (c) Verbal Fluency (VF) Test, which was used to examine language ability, semantic memory, and executive function [35]. And clinical assessment included: (a) Post-concussive symptoms were evaluated based on Rivermead Post-Concussion Symptom Questionnaire (RPCS) [36]; (b) Insomnia Severity Index (ISI) [37]; (c) Posttraumatic stress disorder (PTSD) Checklist-Civilian Version (PCL-C) [38].

### Data Preprocessing

Preprocessing steps of rs-fMRI and T1 data were carried out using the DPABI (http://rfmri.org/dpabi) [39] and SPM (https://www.fil.ion.ucl.ac.uk/spm). Fig. 1 demonstrated the preprocessing steps. In detail, the rs-fMRI data were 1) discarded the first five volumes, 2) slice-time corrected, 3) realigned, 4) regressed out linear trend signal, 24 head motion parameters, and the mean cerebrospinal fluid (CSF) signals, 5) performed temporal scrubbing using motion “spike” (framewise displacement (FD) > 0.5) as separate repressors, 6) filtered by using a band-pass filter (0.01 to 0.15 Hz) for reducing the non-neuronal contribution to BOLD signals which is in line with prior WM FC studies [27,29], 7) spatial smoothed (full width at half maximum = 4 mm) of WM and GM signals separately within individual WM mask or GM mask to avoid the mixture of WM and GM signals, 8) normalized to standard EPI template and resampled voxel size into 3 mm × 3 mm × 3 mm. For T1 image, the T1 data were segmented into WM, GM, and CSF using SPM’s New Segment algorithm. The T1 segmentation results from individual subjects were co-registered to the functional data space for each individual for further the identification WM or GM mask (the segmentation threshold was set to 0.5 using SPM’s tissue segmentation).

**Fig. 1.**
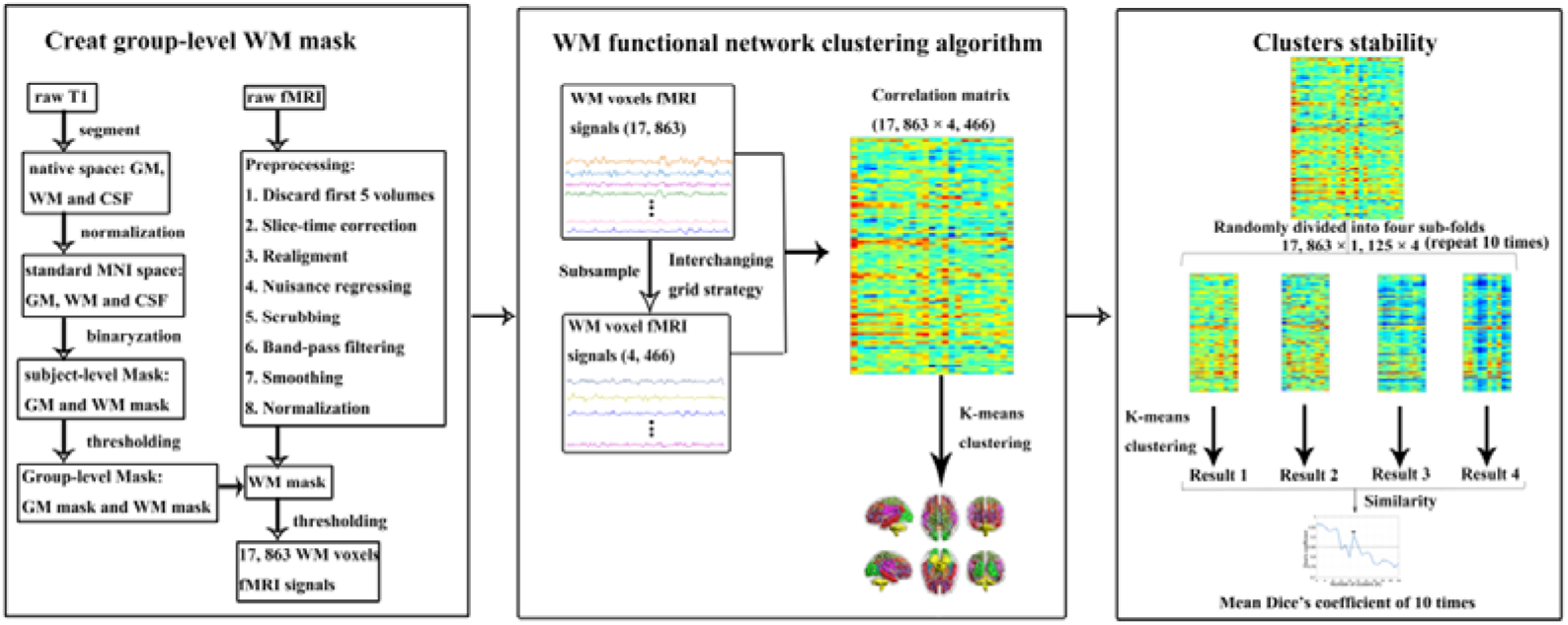
Method overview of white matter functional networks clustering analysis

Preprocessing of DTI data was performed using FSL (https://fsl.fmrib.ox.ac.uk/fsl) software [40], the preprocessing steps included skull stripping of the b0 image, eddy current correction, head motion correction, and registering and aligning to T1 image by mutual information algorithm. The DTI and anatomical T1 images were normalized into MNI space.

### Clustering WM-FNs

The processed resting-state data of mTBI patients in the acute stage and HCs subjects were used to cluster WM-FNs.

First, the segmentation results about the T1 image of each individual were used to create the group WM and GM mask (Fig. 1). For each voxel of each subject, the identification of WM, GM, or CSF was depended on the maximum probability value of the T1 segmentation results. The WM, GM, and CSF mask for each individual were generated subsequently. And the percentage of participants classified as WM or GM was obtained by averaging across all participants. For WM, voxel with a percentage greater than 60% of subjects was identified as a group-level WM mask. For GM, a loose threshold of a percentage greater than 20% of subjects was identified as a group-level GM mask but excluded the voxels included in the WM mask. Of note, to correctly classify deep brain structures [41], the subcortical areas (according to the Harvard-Oxford Atlas [42]) were excluded from the WM mask. Besides, the resulting masks were used to compare with the preprocessed functional data, then we excluded voxels that were identified as GM or WM but without functional data in >20% of the subjects. The group-level WM mask was further co-registered to the functional data space and resampled for rs-fMRI processing. The group-level WM mask contained 17863 voxels.

After generating the GM and WM mask, the K-means clustering algorithm on the averaged group-level resting-state correlation matrices was used to obtain WM-FNs (Fig. 1). To reduce computational complexity, the 17863 voxels of the WM mask were subsampled into 4466 nodes using an interchanging grid strategy [43]. Specifically, any second voxel was extracted along the rows and columns and then moved one between the two slices. The Pearson correlation matrix was calculated between all WM voxels and sub-sample nodes to obtain the correlation matrix (17863 × 4466) for each subject. K-means clustering method (distance metric-correlation, 10 replicates) from machine learning was carried out in the mean correlation matrices. Given clustering method was carried out for an unequal number between patients and the HCs group, the clustering results have the possibility to focus more on the patients, thus lead to the results more inclined to patients. Therefore, the correlation matrix was first averaged in HCs group and mTBI group separately, and then averaged again to obtain final correlation matrix which used for clustering.

Furthermore, to identify the most reliable number of WM-FNs, the stability of each number of clusters ranging from 2 to 22 was evaluated. The whole connectivity matrix (17863 × 4466) was randomly divided into four sub-folds (17863 × 1125). For each kinds of cluster number, each sub-fold carried out the same clustering computation separately so that four clustering results were obtained. To evaluate the similarity of clustering between different sub-folds, the adjacency matrix for each sub-fold was calculated. Then these adjacency matrices were compared using Dice’s coefficient. The stability of the number of clusters was evaluated according to the averaged coefficient of Dice. Furthermore, to ensure the stability of the above results, the above process was repeated for 10 times. Afterward, the average Dice’s coefficient value of the 10 times was taken as the final Dice’s coefficient.

The WM-FNs atlas at the acute stage was also used to analyze the pathological changes of WM-FNs in the chronic stage. To avoid the alteration of the results by the functional reconstruction of WM-FNs in the chronic stage, the WM-FNs clustering of the chronic stage were added (see Supplementary material).

### Similarity between WM-FNs and DTI fiber tracts

The spatial similarity between the WM-FN and DTI fiber tracts was evaluated to investigate the spatial correspondence between them. To identify DTI tracts, the automated fiber quantification algorithm was used to automatically identify 20 fiber tracts depended on the JHU WM tractography atlas. When the voxels where a DTI fiber tract identified in > 2 subjects and belonged to the group-level WM mask constructed previously, the voxels were incorporated into the DTI fiber tract. It should be noted that symmetrical DTI fiber tracts in two hemispheres of the brain were combined as one tract. Furthermore, to investigate the similarity between WM-FNs and DTI fiber tracts, the number of overlapping voxels of each WM-FN and each DTI fiber tract were calculated. The percentage of voxels in the WM-FN which can be identified as part of the DTI tracts (overlap/WM-FN) was also calculated.

### Functional connectivity

To evaluate the interaction within the resulting WM-FNs, the average signal time-courses from WM-FNs were extracted by averaging the signal values across all voxels which were belonged to each network for each individual. The Pearson’s correlation between any two WM-FNs’ time-courses for each individual was computed. Besides, to evaluate the interaction between the WM-FNs and GM-FNs, the GM-FNs atlas which was generated by the same clustering procedure [44] was used, and Pearson’s correlation between the WM-FNs and GM-FNs was also computed. These correlation coefficients were averaged across subjects to obtain a group level matrix representing the relationship between WM and GM-FN. Of note, the Pearson’s correlation coefficients were transformed into the Fisher z score before statistical analysis.

### Statistical analysis

The normality distribution of all cognitive and clinical continuous variables was tested by the Shapiro-Wilk W test. The group differences about cognitive and clinical continuous variables were calculated by the independent two-sample t-test and Mann-Whitney test based on data normality, respectively. Chi-square was used to compare cognitive and clinical categorical variables, with a significant set at p < 0.05.

And independent two-sample t-test was used to show the differences between the mTBI and HCs about the z-score of FC (Pearson’s correlation coefficient), with a significance set at p < 0.05 (false discovery rate corrected, FDR corrected). Since the age of chronic stage patients and HCs was not very well matched (p = 0.064), age was regressed in the statistical analysis of the chronic stage to avoid the influence of brain development.

### Correlation between altered WM-FNs and cognitive and clinical assessments

To evaluate the relationship between altered imaging results and cognitive and clinical assessments, we calculated the Spearman correlation between altered FC and cognitive and clinical variables since the cognitive and clinical variables were not normally distributed.

## Results

### Demographic, cognitive and clinical characteristics

Sixteen patients with mTBI in the acute stage and four HCs with head motion scans exceeding 2 mm and/or 2° rotation were excluded. The final analysis included 97 mTBI patients in the acute stage, 56 mTBI patients in the chronic stage, and 43 HCs. The mTBI patients did not differ from HCs regarding age, education, gender, and mean FD (Table 1).

**Table 1.** Demographic characteristics of the mTBI patients and HCs

| characteristics | mTBI (n=97)<br>(acute stage) | mTBI (n=56)<br>(chronic stage) | HCs (n=43) | p1 | p2 |
| --- | --- | --- | --- | --- | --- |
| Age (years) | 38.99 ± 13.76 | 35.32±15.32 | 39.93 ± 13.09 | 0.454 | 0.064 |
| Gender (male:female) | 48:49 | 33:23 | 20:23 | 0.745 | 0.220 |
| Education (years) | 7.95 ± 3.99 | 8.54±3.77 | 8.77 ± 5.50 | 0.598 | 0.804 |
| Mean FD | 0.11±0.10 | 0.11±0.07 | 0.11±0.06 | 0.930 | 0.801 |
| **Cognitive and Clinical assessments** |  |  |  |  |  |
| TMT-A | 73.07±50.55 | 44.86±35.88 | 52.30±29.97 | 0.023 | 0.109 |
| DSC | 31.99±16.30 | 44.54±15.34 | 39.63±17.72 | 0.021 | 0.283 |
| Forward DS | 7.75±1.56 | 8.57±1.59 | 7.81±1.60 | 0.835 | 0.037 |
| Backward DS | 3.80±1.47 | 4.79±1.60 | 3.95±1.45 | 0.564 | 0.008 |
| VF | 16.43±5.47 | 18.86±6.19 | 17.28±6.02 | 0.389 | 0.147 |
| RPCS | 11±8.13 | 5±5.30 | 2.49±2.88 | <0.001 | 0.014 |
| ISI | 7.08±5.79 | 3.39±4.53 | 1.44±2.64 | <0.001 | 0.009 |
| PCL-C | 25.12±6.18 | 21.66±5.69 | 17±0 | <0.001 | <0.001 |

Abbreviations: mTBI, mild traumatic brain injury; HCs, healthy controls; TMT-A, trail making test A; DSC, digit symbol coding; DS, digit span; VF, verbal fluency; RPCS, rivermead post-concussion symptom questionnaire; ISI, insomnia severity index; PCL-C, posttraumatic stress disorder checklist-civilian version. p1 represents the p-values from statistical test between mTBI in acute stage and HCs. p2 represents the p-values from statistical test between mTBI in chronic stage and HCs.

Significant differences were revealed between patients with mTBI in acute stage and HCs in TMT-A (p = 0.023), DSC (p = 0.021), RPCS (p < 0.001), ISI (p < 0.001), and PCL-C (p < 0.001) (Table 1, p1 represent the p-value from statistical test between acute mTBI patients and HCs). And significant differences in Forward DS (p = 0.037), Backward DS (p = 0.008), RPCS (p = 0.014), ISI (p = 0.009), and PCL-C (p < 0.001) were observed between patients with mTBI in chronic stage and HCs (Table 1, p2 represent the p-value from statistical test between chronic mTBI patients and HCs). The detailed demographic information and cognitive and clinical assessments of mTBI and HCs were demonstrated in Table 1.

### WM-FNs

The coefficient of Dice was adopted to evaluate the stability of WM-FNs number, the results demonstrated the most stable (Dice’s coefficient > 0.85) with the largest number of WM-FNs was 11 (Fig. 2). The WM-FN is given a putative network name based on the anatomical location. The WM-FNs can be named as: WM1 (IFOF network), WM2 (corona radiate network), WM3(anterior temporal network), WM4 (orbito-frontal network), WM5 (pre/postcentral network), WM6 (superior longitudinal fasciculus network), WM7 (ventral frontal network), WM8 (temporoparietal network), WM9 (Rolandic network), WM10 (cerebellar network), WM11 (occipital network) (Fig. 2, Table 2).

**Fig. 2.**
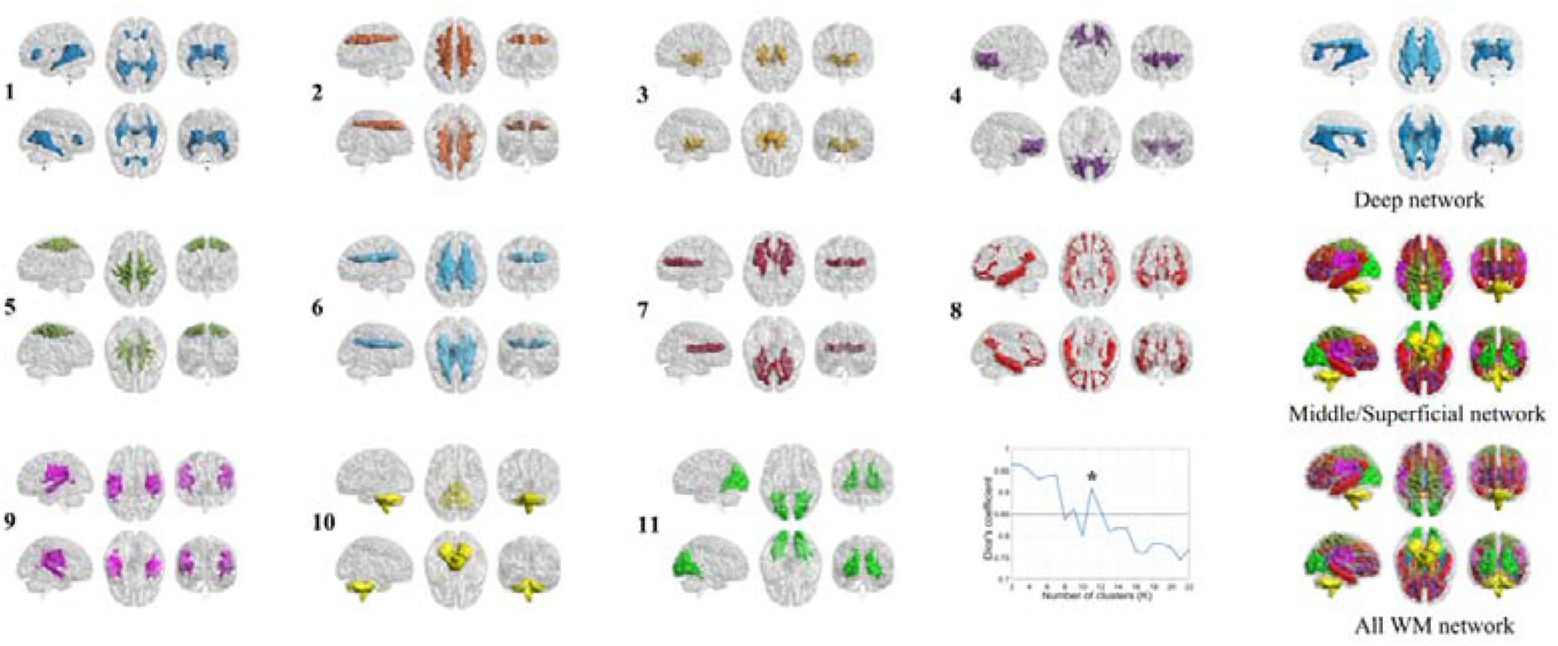
Clusters stability, white matter functional networks (WM-FNs) of most suitable clusters (k = 11), and middle/superficial and deep WM-FNs

**Table 2.**
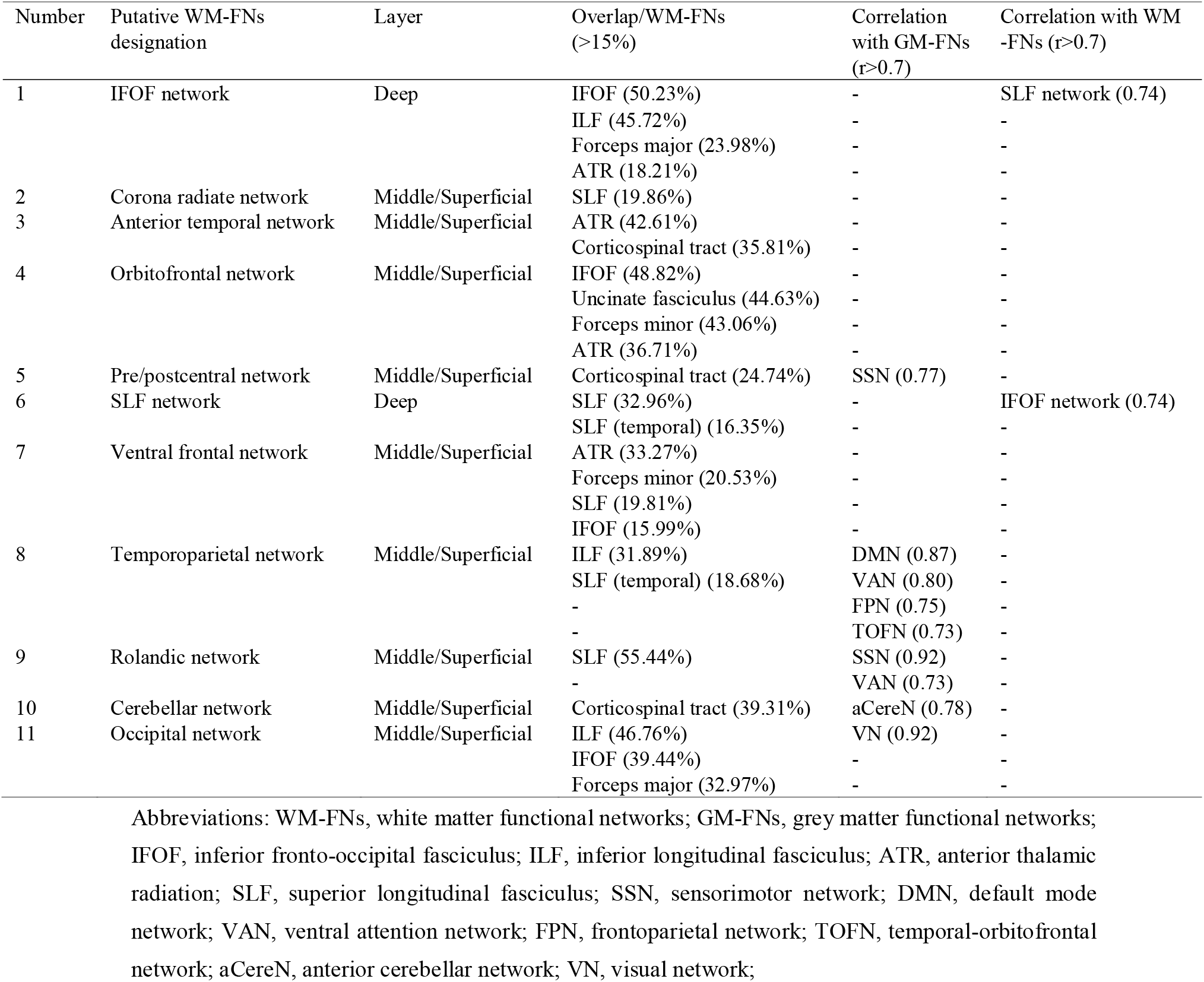
Overlap of WM-FNs with white matter fiber tracts, correlation with GM-FNs and WM-FNs

Besides, based on the spatial distribution [24], we defined middle/superficial WM-FNs (WM-FN 2, 3, 4, 5, 7, 8, 9, 10, 11) and deep WM-FNs (WM-FN 1, 6) (Fig. 2, Table 2)

### Similarity between WM-FNs and DTI fiber tracts

Four WM-FN (2, 5, 9, 10) of the 11 networks observed spatial correspondence with specific DTI anatomical tracts (e.g., network 9 and superior longitudinal fasciculus;) (Fig. 3, Table 2). And the other networks spatial corresponded to multiple DTI tracts (e.g., network 1 spatial corresponded to both IFOF and inferior longitudinal fasciculus;) (Fig. 3, Table 2).

**Fig. 3.**
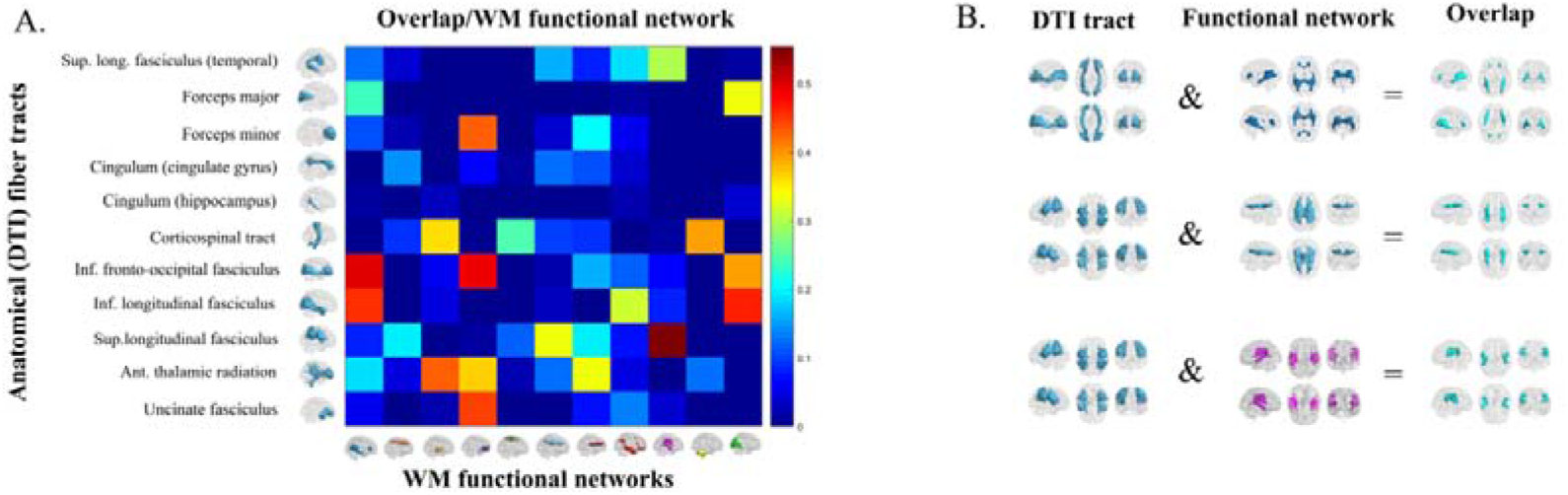
The spatial similarity between the white matter functional networks (WM-FNs) and diffusion tensor imaging (DTI) fiber tracts. (a) the ratio between the voxel number of overlapping and the voxel number of WM-FN. (b) Examples about the spatial interaction between the WM-FN and DTI fiber tract

### Functional connectivity

#### Functional connectivity and its association with cognitive deficits in the acute stage

Across subjects, middle/superficial WM-FNs demonstrated high correlation in their spontaneous activity with their adjacent GM-FNs and some distributed GM-FNs, such as occipital network and visual network (r = 0.92), rolandic network and sensorimotor network (r = 0.92), temporoparietal network and default mode network (r = 0.87) (Fig. 4 (a), Table 2). Besides, a relatively weak correlation (r < 0.55) was revealed between the deep WM-FNs and GM-FNs (Fig. 4 (a)).

**Fig. 4.**
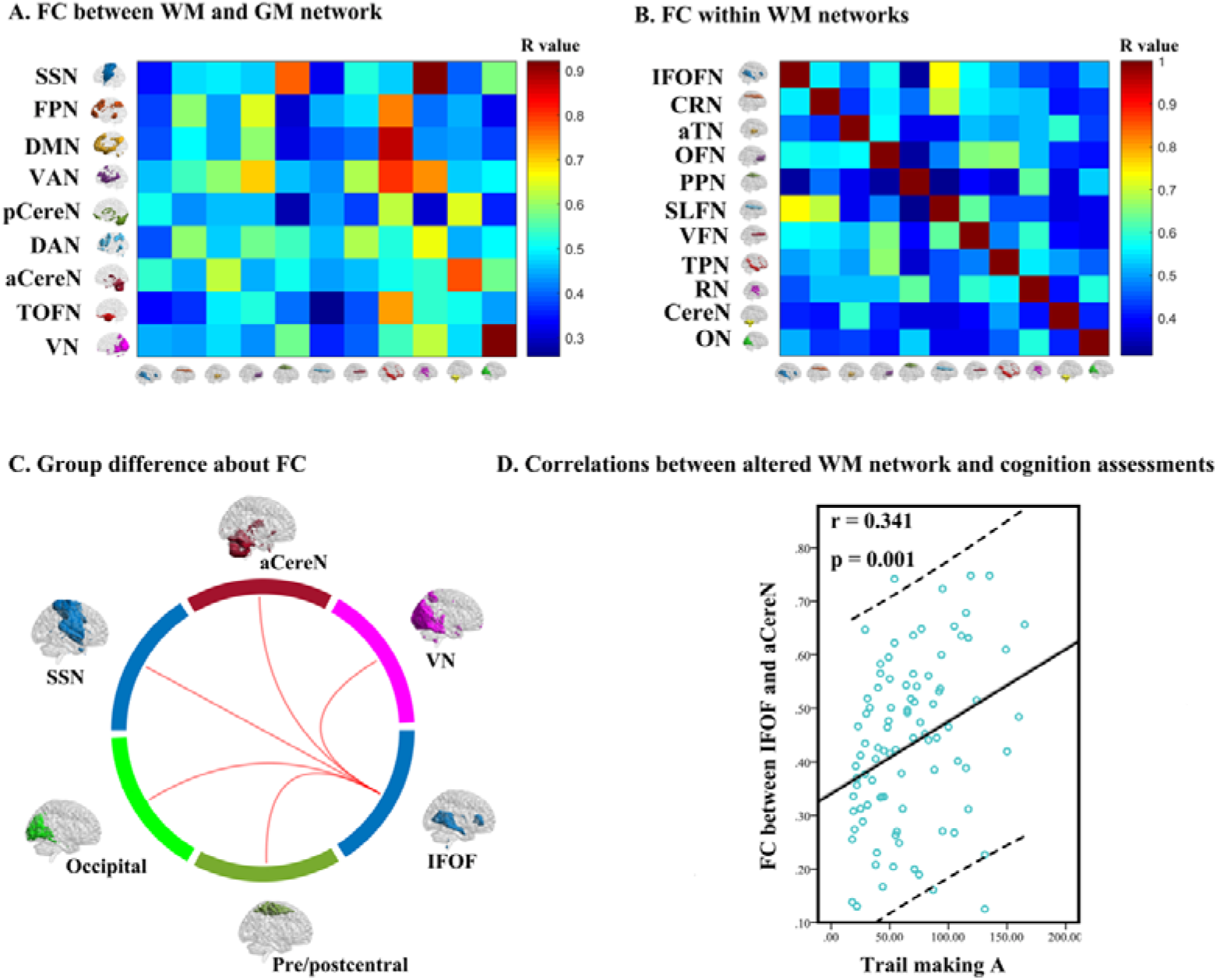
Functional connectivity (FC) of white matter functional networks (WM-FNs) in the acute stage mild traumatic brain injury (mTBI) patients and healthy controls (HCs). (a) Average FC between the WM-FNs and grey matter functional networks (GM-FNs). (b) Average FC within WM-FNs. (c) Group differences between patients and HCs in FC within WM-FNs or between WM-FN and GM-FN. The red connections represented significantly increased FC in mTBI patients when compared with HCs (FDR corrected, p < 0.05). (d) Correlations between abnormal FC and cognition assessments

Across subjects, between adjacent WM-FNs, such as the superior longitudinal fasciculus network and IFOF network, showed a strong correlation (r = 0.74) (Fig. 4 (b), Table 2). Between distant WM-FNs, such as cerebellar WM-FN and the other WM-FNs, showed relatively weak correlation (r < 0.60) (Fig. 4 (b)).

Compared to HCs, patients with mTBI showed increased FC between the deep WM-FN and low-level primary sensorimotor cortical GM-FNs. In detail, mTBI patients showed increased FC between IFOF WM-FN and visual GM-FN, IFOF WM-FN and anterior cerebellar GM-FN, IFOF WM-FN and sensorimotor GM-FN (Fig. 4 (c), Table 3).

**Table 3.** Group differences between mTBI patients in the acute stage and HCs in FC within WM-FNs and between WM-FNs and GM-FNs

| WM-FNs | Functional networks | t value | p value |
| --- | --- | --- | --- |
| Inferior fronto-occipital fasciculus WM-FN | Visual GM-FN | 4.085 | 0.007 |
|  | Anterior cerebellar GM-FN | 3.266 | 0.048 |
|  | Sensorimotor GM-FN | 3.252 | 0.048 |
|  | Occipital WM-FN | 3.546 | 0.030 |
|  | Pre/postcentral WM-FN | 3.205 | 0.046 |

Abbreviations: mTBI, mild traumatic brain injury; HCs, healthy controls; FC, functional connectivity; WM-FNs, white matter functional networks; GM-FNs, grey matter functional networks; WM-FN, white matter functional network; GM-FNs, grey matter functional network; Note: FDR corrected $p < 0.05$ .

Compared to HCs, patients with mTBI also showed increased FC between deep WM-FN and low-level primary sensorimotor superficial WM-FN. In detail, patients with mTBI showed increased FC between IFOF WM-FN and occipital WM-FN. Also, patients with mTBI showed increased FC between IFOF WM-FN and pre/postcentral WM-FN (Fig. 4 (c), Table 3).

Taken together, the IFOF WM-FN is the most affected WM-FN, which showed increased connectivity with low-level primary sensorimotor functional networks in acute mTBI patients.

The FC between IFOF WM-FN and anterior cerebellar GM-FN was positively correlated with TMT-A score (Fig. 4 (d)).

#### Functional connectivity and its association with cognitive deficits in the chronic stage

Similar to the acute stage, the interaction between WM-FNs and GM-FNs in the chronic stage showed a relatively high correlation between middle/superficial WM-FNs and adjacent GM-FNs, such as the occipital network and visual network (r = 0.92), rolandic network and sensorimotor network (r = 0.92), temporoparietal network and default mode network (r = 0.87). The deep WM-FN and GM-FNs showed a relatively weak correlation (r < 0.55) (Fig. 5 (a)).

**Fig. 5.**
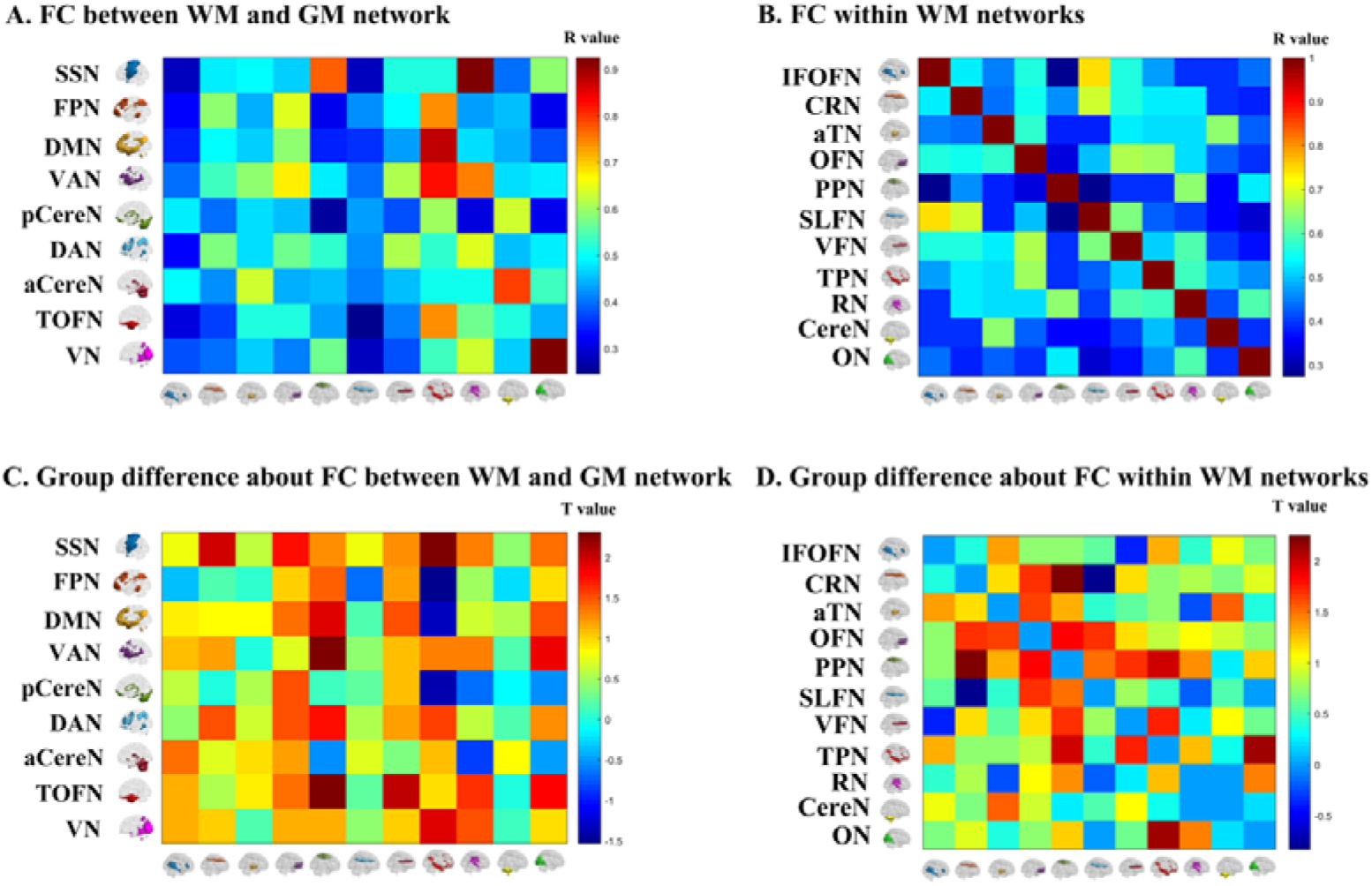
Functional connectivity (FC) of white matter functional networks (WM-FNs) in the chronic stage mild traumatic brain injury (mTBI) patients and healthy controls (HCs). (a) Average FC between the WM-FNs and grey matter functional networks (GM-FNs). (b) Average FC within WM-FNs. (c) Group differences of FC within distinct WM-FNs between mTBI patients and HCs. (d) Group differences of FC between WM-FNs and GM-FNs between mTBI patients and HCs. The color bar in (a) and (b) shows the Pearson’s correlation coefficient (R value). The color bar in (c) and (d) shows the T value from the two-sample t-test

The interaction between pair of WM-FNs in the chronic stage was also similar to the acute stage with a strong correlation between adjacent WM-FNs, such as the superior longitudinal fasciculus network and IFOF network (r = 0.75) (Fig. 5 (b)). A relatively weak correlation was presented between distant WM-FNs, such as cerebellar WM-FN and the other WM-FNs (r < 0.60) (Fig. 5 (b)).

The mTBI patients did not differ from HCs regarding FC between WM and GM-FNs (Fig. 5 (c)). Also, mTBI patients and HCs did not differ regarding FC within WM-FNs (Fig. 5 (d)).

Besides, even the WM-FNs were constructed by the patients in chronic stage and HCs, the results remain stable (see in supplementary material).

## Discussion

The present study, for the first time, investigated the dynamic change of WM-FN in mTBI within a longitudinal study. Eleven distinct WM-FNs were identified by clustering voxel-wise WM functional connectivity. These networks are corresponded to either a specific or multiple anatomical DTI WM tracts. The altered FC of WM-FN was observed in the acute stage but not in the chronic stage, suggesting a possible compensatory mechanism in the chronic recovery stage. Importantly, the IFOF WM-FN demonstrated enhanced FC both with the low-level GM primary sensorimotor networks, and with the low-level WM primary sensorimotor networks in acute mTBI patients. These enhanced FC may represent a compensatory or protective mechanism in response to cognitive deficits in acute mTBI. The convergent damage of IFOF network highlighted the importance of IFOF WM-FN in our understanding of the pathophysiology mechanism of mTBI patients, which can further aid the promotion of new treatment intervention that targeting IFOF network in the acute stage.

### The spatial similarity between WM-FNs and DTI fiber tracts

In previous studies, the WM tracts integrity analysis of mTBI patients were observed mainly based on structural MRI, such as DTI. The current study reported a novel observation of functional communication of WM in mTBI patients. The constructed WM-FNs were corresponded to either a specific or multiple anatomically DTI WM tracts. This is consistent with the previous study [27] to support the notion that multiple DTI tracts collaborate with each other to guarantee the specific brain function. The analysis of spatial interacting patterns between DTI fiber tracts and WM-FNs helps to clarify which DTI fiber tract can be reflected by the WM-FN, especially by the deep WM networks. For example, the identified IFOF functional network was mainly a spatial interaction with IFOF fiber tract (Fig. 3, Table 2). Thus, the IFOF functional network was assumed to mainly reflect the function of IFOF fiber tract. It should be noted that it is difficult to describe middle/superficial WM-FNs using DTI fiber tract since they combine many small fibers with different orientations [45]. Therefore, the function of middle/superficial WM-FNs is mainly reflected by their connectivity with cortical GM-FNs. For instance, the occipital WM-FN showed strong connectivity with cortical visual GM-FN, so that the occipital WM-FN was assumed to also reflect the visual function to some extent. In sum, and together with previous study [46,21,27], these results provide further support that WM-FNs have corresponding structural base.

### Functional connectivity in the acute stage

Intriguingly, the IFOF WM-FN was found to be the most affected deep WM-FN in mTBI patients. The IFOF WM-FN showed enhanced FC both with the low-level GM primary sensorimotor networks (visual network, anterior cerebellar network, and sensorimotor network), and with the low-level WM primary sensorimotor networks (occipital network and pre/postcentral network) in patients with mTBI at the acute stage. Previous studies have reported the damaged structure property in IFOF in patients with mTBI [47,48]. Here, for the first time, the present study showed that increased IFOF FC with low-level WM and GM primary sensorimotor networks in patients with mTBI. In general, the increased positive FC reflects an increased integrative ability sub-serving similar goals [49]. Hence, the increased IFOF FC may reflect a compensatory mechanism in mTBI. As a long-distance associative tract, the IFOF plays a crucial role in connecting the occipital lobe, the parietal lobe, and the poster-temporal cortex with the frontal lobe [13], which was particularly vulnerable to the mechanical trauma of TBI [11]. Correspondingly, the IFOF is crucial in sending the processed visual information from the occipital lobe to frontal lobe, where information was further translated into semantic information [50]. The observed increased connectivity between IFOF FC with low-level WM and GM primary sensorimotor networks may represent a compensatory effect in response to cognitive deficits. These enhanced FCs may indicate mTBI patients employed higher cognitive effort in integrating dynamic environmental sensory information and high-order cognitive function to coordinate their behaviors and thoughts to fit with contextual information. This interpretation is supported by previous studies which demonstrated abnormal activation of the intrinsic rich-club network in mTBI patients [51] and greater network strength in mTBI patients without post-concussive syndrome [52]. Both studies suggested a potential compensatory or protective mechanism in mTBI patients. The current findings highlighted the importance of the IFOF WM-FN in such a compensatory mechanism, in which IFOF WM-FN may excessively contribute to integrating sensory information from GM and WM primary sensory-related network with high-order cognitive information in acute mTBI patients. The most affected damage in mTBI was located in IFOF network, which implies the crucial role of IFOF network in our understanding of the pathophysiology mechanism of mTBI patients. This observation can further aid the promotion of new treatment intervention targeting IFOF network in the acute stage.

### The association between altered WM-FNs and cognitive assessments

Besides, the FC between IFOF WM-FN and anterior cerebellar GM-FN was positively correlated with TMT-A score in mTBI patients at acute stage. This correlation suggested a deficit of the information processing speed in acute mTBI patients. Information processing speed is depend on large-scale and long-distance neural network communications, which is among the earliest and most prominent cognitive manifestations in mTBI [53,54]. Thus, the deficit of information processing speed was assumed to be caused by the damage of IFOF network in acute mTBI patients to some extent.

### Functional connectivity in the chronic stage

Furthermore, mTBI patients at the chronic stage did not differ from HCs in FC within WM-FNs or between GM-FN and WM-FN. A previous meta-analysis indicated that the damage effects for memory and fluency were greatest in the acute stage (less than 3 months postinjury) and no residual impairment by 3 months postinjury [55]. Besides, a recent large sample longitudinal neuroimaging study of TBI revealed a ‘U-shaped’ curve in sub-acute, 1 year, and 5 years postinjury in the number of fractional anisotropy abnormalities [56]. Furthermore, a previous study demonstrated a functional and structural abnormal connectivity in early phase of mTBI, but a considerable compensation of functional and structural connectivity subsequent to the acute phase [57]. Combining previous studies and the current findings, it is evident that the compensatory mechanism of brain function contributed to the recovery of cognition deficits in mTBI patients at the chronic stage.

### Limitation

Notwithstanding its implication, limitations of the current study should be acknowledged and addressed. Some researchers have speculated that the WM signals might have infiltrated from the GM for partial volume effect. To avoid the influence of GM signals, the spatial smooth of WM and GM signals were separately carried out within individual WM mask or GM mask to avoid the mixture of WM and GM signals. Besides, only the data of mTBI patients in the acute stage and the chronic stage (6-12 months postinjury) was collected, the data for longer follow-up, such as 5 years postinjury should be collected as a follow-up. Based on a previous study [56], the significant abnormalities of FA in TBI were observed at the sub-acute and 5 years postinjury, but not 1-year postinjury. Therefore, it is necessary to follow patients for a longer period to explore the development of the observed deficits of WM-FNs.

## Conclusion

In summary, this study constructed WM-FNs based k-means clustering algorithm and revealed an excessive interaction between deep WM-FNs and low-level primary sensorimotor networks, which can be regarded as a compensatory mechanism of cognitive deficits in mTBI patients in the acute stage. Of note, patients with mTBI did not differ from HCs at the chronic stage in FC within WM-FNs or between GM and WM-FN, suggesting a possible compensatory mechanism of brain function contributed to the recovery of cognition deficits in mTBI patients at the chronic stage. The convergent damage of IFOF network highlighted its key role in our understanding of the pathophysiology mechanism of mTBI patients and thus might be regarded as a biomarker in the acute stage and a potential indicator of treatment effect.

## Supplementary material

### Detailed inclusion and exclusion criteria of mTBI patients

In detail, patients with mTBI were selected with the following criteria: (a) Glasgow Coma Score of 13-15; (b) one or more of the following: confusion or disorientation, post-traumatic amnesia for less than 24 hours, loss of consciousness for 30 minutes or less, and/or other transient neurological abnormalities such as focal signs, seizure, and intracranial lesion not requiring surgery; (c) diagnosed within 1 week after onset of mTBI. The exclusion criteria included: (a) a history of a previous brain injury, neurological disease, long-term psychiatric history, or a history of concurrent substance or alcohol abuse; (b) a structural abnormality in neuroimaging (CT and MRI); (c) intubation and/or skull fracture, and administration of sedatives; (d) the manifestation of mTBI caused by medications, alcohol, drugs for other injuries (such as systemic injuries, facial injuries, or intubation); (e) other problems (such as psychological trauma, language impairment, or coexisting medical conditions); (f) caused by penetrating craniocerebral injury.

### Image acquisition

The rs-fMRI data was acquired using a gradient-recalled echo planar imaging (EPI) sequence, and the scan parameters were as follows: repetition time (TR) = 2500 ms, echo time (TE) = 30 ms, slice thickness = 3 mm, flip angle (FA) = 90°, field of view (FOV) = 216 mm × 216 mm, matrix size = 64 × 64. One hundred and eighty volumes were obtained. The scan parameters of High-resolution T1-weighted 3D BRAVO sequence were as follows: TR = 8.15 ms, TE = 3.17 ms, slice thickness = 1 mm, FA = 9°, FOV = 256 mm × 256 mm. The scan parameters of DTI were as follows: TR = 8000 ms, TE = 68ms, FA = 90°, slice thickness = 2mm, slices = 75, matrix size = 128 × 128, two averages, FOV = 256 mm × 256 mm, voxel size = 2 mm × 2 mm × 2 mm. DTI (b = 1000 s/mm2) used 30 diffusion gradient orientations and the unweighted diffusion imaging (b = 0) repeated five times. During scanning, all participants were instructed to relax, close their eyes, keep awake, and try not to think of anything in particular.

### WM-FNs clustering of the chronic stage

The WM-FNs clustering based on the data of patients in the chronic stage and HCs data were also calculated to avoid the functional reconstruction of WM-FNs in the chronic stage. According to the results of Dice’s coefficient (Supp Fig. 1), we identified 12 WM-FNs (Supp Fig. 1). Compared with HCs, there also was no significant difference between patients and healthy control in functional connectivity between WM-FNs and GM functional connectivity, and no significant difference between patients and healthy control in functional connectivity within WM-FNs in the chronic stage of mTBI (Supp Fig. 2).

---

**Supp Fig. 1.**
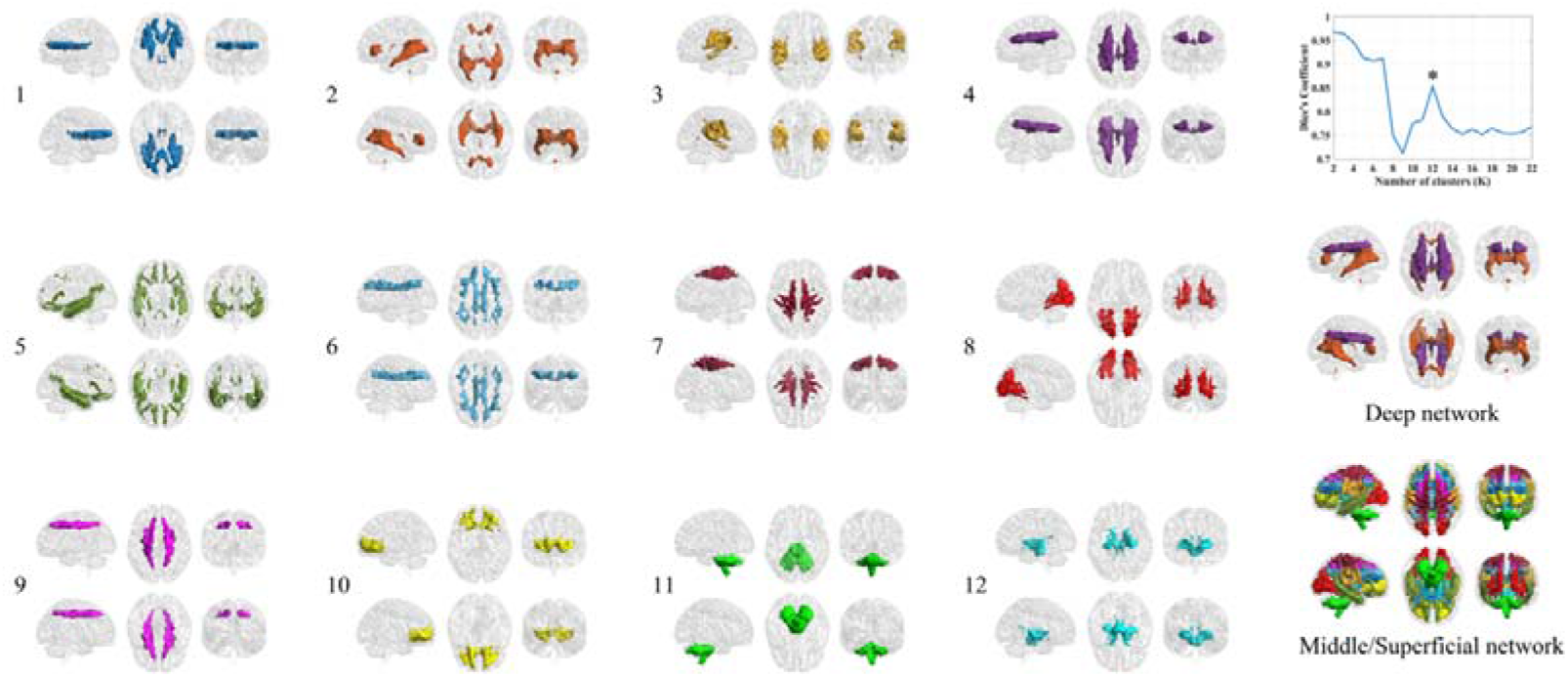
White matter functional networks identified by patients in the chronic stage and healthy control

**Supp Fig. 2.**
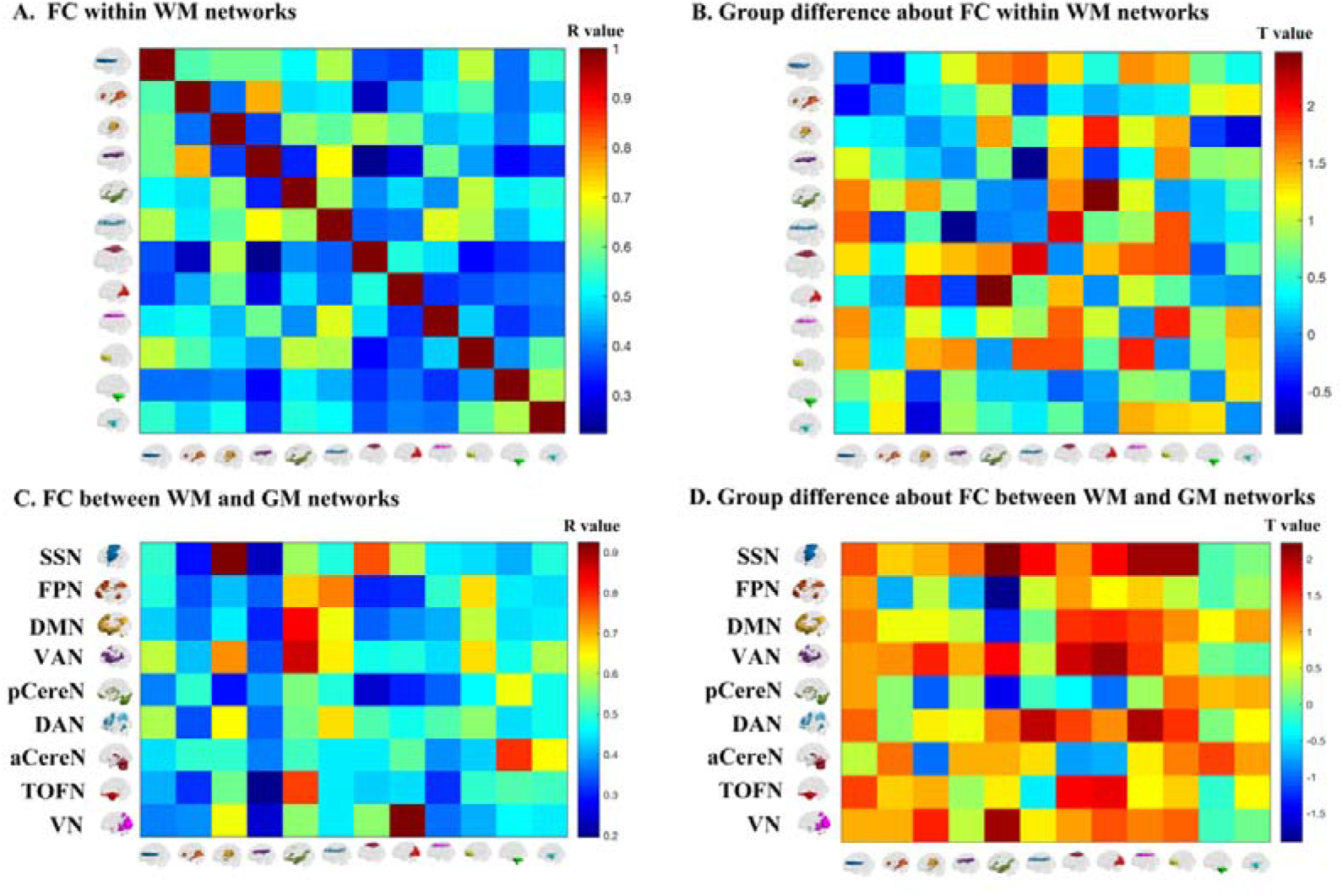
Functional connectivity (FC) of white matter functional networks (WM-FNs) in chronic stage mild traumatic brain injury (mTBI) patients and healthy controls (HCs). (a) Average FC within WM-FNs. (b) Group differences between mTBI patients and HCs in FC within WM-FNs (c) Average FC between the WM-FNs and grey matter functional networks (GM-FNs). (d) Group differences between mTBI patients and HCs in FC between WM-FNs and GM-FNs. The color bar in (a) and (c) shows the Pearson’s correlation coefficient (R value). The color bar in (b) and (d) shows the T value from the two-sample t-test

